# Direct Evidence for Glucose Consumption Acceleration by Carbonates in Cultured Cells

**DOI:** 10.1101/551259

**Authors:** Kenji Sorimachi

**Affiliations:** Educational Support Center, Dokkyo Medical University, Miub, Tochigi 320-0293, Japan; Research Laboratory, Gunma Agriculture and Forest Development, Takasaki, Gunma 370-0851, Japan

## Abstract

Established Py-3Y1-S2 rat fibroblast cells were used to evaluate whether NaHCO_3_ or Na_2_CO_3_ influences glucose metabolism *in vitro*, because factors that contribute to metabolic pathways are much simpler to evaluate in cultured cells than in whole animal bodies. The effects of the carbonates on glucose consumption decreased at high concentrations, >5 mg/ml for Na_2_CO_3_ and >7 mg/ml for NaHCO_3_, because of the increased pH of the culture medium. The effects of the carbonates on glucose consumption were additive with those of vanadium and concanavalin A. Streptozotocin, alloxan, and nicotinamide, which induce diabetes in animals, reduced glucose consumption by Py-3Y1-S2 cells, and the inhibitory effects of these reagents were abolished by both Na_2_CO_3_ and NaHCO_3_. Finally, the carbonates increased lactate production from glucose in the cells, followed by acceleration of lactate secretion into the culture medium. The present study clarified that NaHCO_3_ and Na_2_CO_3_ directly regulate glucose metabolism.

## INTRODUCTION

The prevalence of diabetes mellitus (DM) is increasing worldwide. The International Diabetes Federation reported that there were 425 million adults with diabetes in the world in 2017 [1]. The symptoms of chronic high blood sugar in DM include frequent urination, increased thirst, and increased hunger. If left untreated, DM can lead to serious complications like cardiovascular disease, stroke, chronic kidney failure, foot ulcers, and eye damage. Insulin via injections is used for treatment of type 1 DM, while insulin and several oral drugs that inhibit glucose production from polysaccharides are used for type 2 DM treatment. However, these drugs only treat the symptoms, and there is no basic remedy for DM. Previous studies have investigated whether insulin plays a role in memory formation *in vivo* [2] and whether insulin contributes to dementia, particularly Alzheimer’s disease [3–5].

Regarding its mode of action as a peptide hormone, insulin binds to its receptor on the plasma membrane, leading to intramolecular phosphorylation within the activated receptor as a tyrosine kinase. The signal transduction proceeds through phosphatidylinositol-3-kinase, protein kinase B, and glucose transporter-4 (GLUT-4), with GLUT-4 plus K^+^ accelerating glucose uptake into cells [6]. Vanadium compounds were reported to exhibit insulin-like activity not only *in vitro* [7,8], but also *in vivo* [9–16], and several vanadium compounds have been investigated for their insulin-like activity [17–21]. The insulin-like effects of vanadates are based on inhibition of protein-tyrosine phosphatase [22]. However, to our knowledge, suitable vanadium compounds have not yet been developed as anti-DM drugs because of the serious cytotoxic effects of high vanadium concentrations. It has been suggested that Mt. Fuji subsoil water filtered through basalt can exhibit insulin-like activity, because the water contains vanadium pentoxide (V_2_O_5_) *in vivo* [23]. Recently, we confirmed that Mt. Fuji subsoil water accelerates glucose consumption *in vitro* using established Py-3Y1-S2 rat fibroblast cells [24] and human primary fibroblasts [25]. Vanadium pentoxide is soluble in the alkaline condition, but its water solubility is quite low (0.7–0.8 g/L). Indeed, the pH value of commercial Mt. Fuji subsoil water (Healthy Vana Water) containing 130 μg/L vanadium was 8.3 [24]. If vanadium-containing water can be prepared by mixing a small amount of Mt. Fuji basalt powder with normal water, the vanadium-containing water could be conveniently used instead of Mt. Fuji subsoil water.

Associations between metabolic acidosis, insulin resistance, and cardiovascular risks have been reported since 1924 [26]. Although correction of metabolic acidosis by nutritional therapy and/or oral administration of sodium bicarbonate is widely applied in chronic kidney disease [27–29], it remains unknown whether metabolic acidosis reduces insulin resistance and/or improves insulin effects on target cells in diabetic subjects. Bellis et al. [30] evaluated whether metabolic acidosis correction by sodium bicarbonate administration can improve peripheral endogen insulin utilization by target organs in diabetic subjects with chronic kidney disease treated with oral antidiabetic drugs. They reported that sodium bicarbonate administration improved insulin resistance in chronic kidney disease at the serum glucose and insulin levels. However, they commented that further investigations were required to evaluate their results in diabetic and non-diabetic chronic kidney disease patients. Indeed, another group reported that sodium bicarbonate treatment did not improve insulin sensitivity and glucose control in non-diabetic older adults [31]. These contradictory findings seem to arise from the different experimental systems investigated with many different factors. The present study was designed to clarify whether carbonates (NaHCO_3_ and Na_2_CO_3_) can directly influence glucose metabolism in simple cultured cells.

## MATERIALS AND METHODS

### Cell culture

Py-3Y1-S2 rat fibroblast cells were cultured in Dulbecco’s minimum essential medium (DMEM) containing 5% fetal bovine serum (Gibco, Thermo Fisher Scientific K.K., Tokyo, Japan) in the presence of 5% CO_2_ at 37°C. Stock cells cultured in 25-cm^2^ plastic culture flasks were trypsinized and inoculated into 24-well culture plates. After the cells had almost reached confluent monolayers, they were cultured in the presence or absence of NaHCO_3_, Na_2_CO_3_, and other chemical compounds like vanadium, concanavalin A (Con A), nicotinamide, streptozotocin (STZ), and alloxan monohydrate.

### Glucose assay

Glucose concentrations in culture media were measured by a previously reported method [24] using a Glucose CII-test (Wako Pure Chemical, Osaka, Japan). For this, a 10–50-μl aliquot of culture medium was mixed with the assay solution (0.5 ml) and the absorbance at 505 nm was measured after 15 min. To express the effects of reagents on glucose consumption, the amount of consumed glucose in the sample medium was divided by that in the control medium (Milli-Q water) without a test reagent.

### Lactate assay

Lactate concentrations in culture media were measured with a Lactate Assay Kit-WST (Dojin Chemicals, Tokyo, Japan) according to the manufacturer’s protocol. Briefly, a 20-μl aliquot of sample solution was mixed with 80 μl of working solution and incubated at 37°C for 30 min. After the enzyme reaction, the absorbance at 450 nm was measured to calibrate the lactate concentration.

### Protein assay of cultured cells

Cellular protein assays were carried out using a previously reported method [24]. Cells cultured in 24-well plates were washed with phosphate-buffered saline and treated with 0.2 ml of Mammalian Protein Extraction Reagent (Thermo Scientific, Rockford, IL, USA). An aliquot of the solubilized cell solution (4 μl) was added to 156 μl of H_2_O, and mixed with Bio-Rad Protein Assay Dye Reagent (Bio-Rad Laboratories, Hercules, CA, USA). The absorbance at 595 nm was measured. Bovine serum albumin was used as a standard.

### Chemicals

DMEM powder (Misui, Tokyo, Japan) was dissolved in Milli-Q water, and autoclaved. The medium (1 L) was neutralized by adding 4 ml of 10% NaHCO_3_, according to the technical protocol recommended by the manufacturer. NaHCO_3_, Na_2_CO_3_, vanadium, nicotinamide, STZ, and alloxan monohydrate were purchased from Wako Pure Chemical Industries Ltd. (Osaka, Japan). Con A was obtained from Sigma Chemical Co. (St. Louis, MO, USA).

### Statistical analysis

Statistical calculations by the *t*-test were performed using Microsoft Excel (version 2010). Values of p < 0.05 and p < 0.01 were considered significant and highly significant, respectively.

## RESULTS AND DISCUSSION

### Effects of NaHCO_3_ and Na_2_CO_3_

To prepare vanadium-containing water, Mt. Fuji basalt powder (1–2-μm diameter) was treated with 1 mg/ml NaHCO_3_, which is safe for the human body, and this water was used to make the cell culture medium. The cell culture medium containing Mt. Fuji basalt-treated water increased glucose consumption in cultured cells (Fig. 1). However, an increase in Mt. Fuji basalt powder for water treatment from 5 mg/10 ml to 200 mg/10 ml did not cause any further increase in glucose consumption by cultured cells. These findings indicate that Mt. Fuji basalt powder treatment may not be involved in acceleration of glucose consumption by cultured cells. Indeed, only NaHCO_3_ or Na_2_CO_3_ accelerated glucose consumption by cultured cells (Fig. 1).

**Figure 1.**
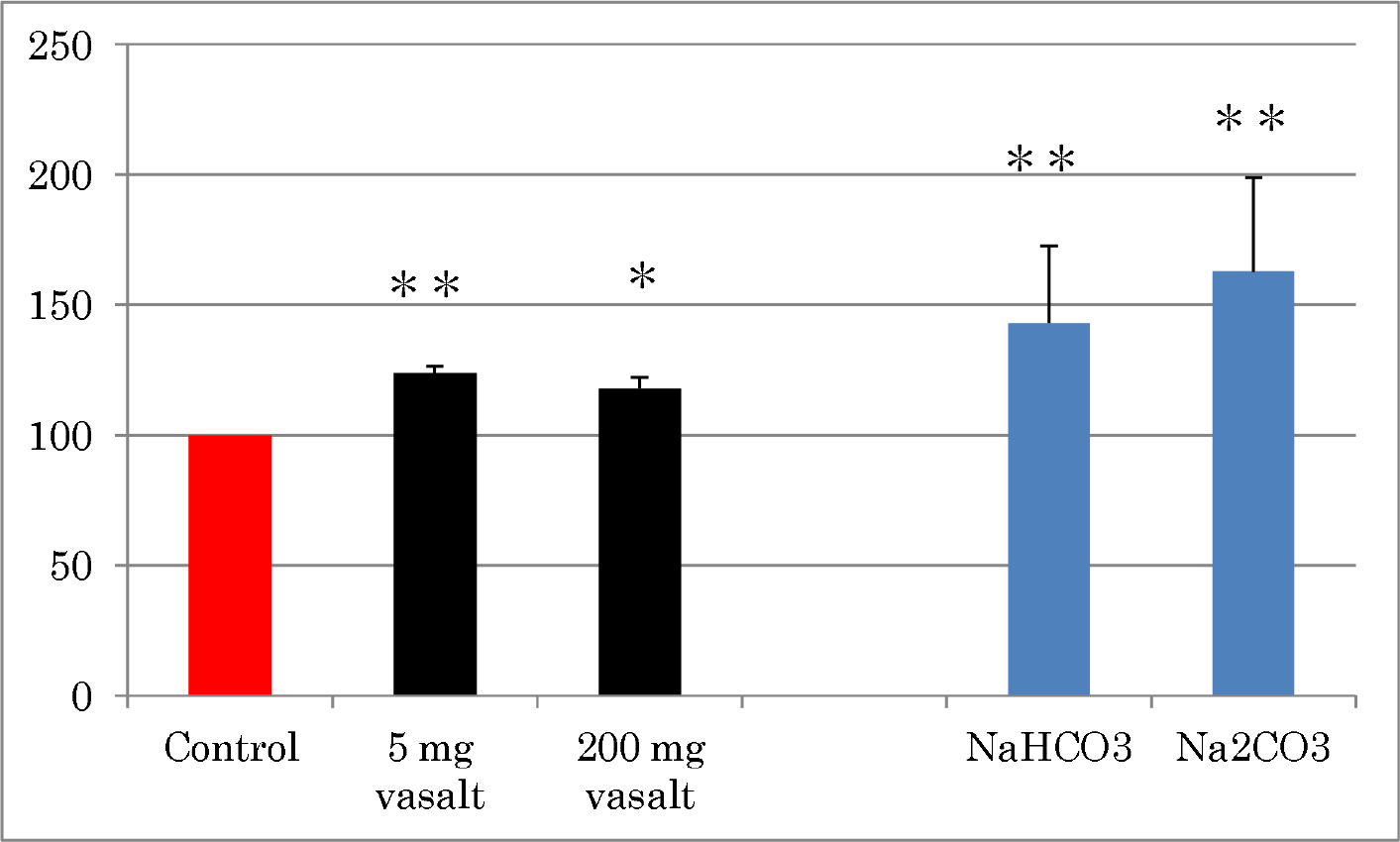
Effects of basalt powder on glucose consumption by cultured Py-3Y1-S2 cells in the presence of NaHCO_3_ or Na_2_CO_3_. Mt. Fuji basalt powder with ~2-μm diameter was kindly supplied by Mt. Fuji Yogan Institute (Yamanashi, Japan), and 5 or 200 mg of powder was treated with 1 mg/ml NaHCO_3_ for 30 min at room temperature. After treatment with NaHCO_3_, the solution was centrifuged at 3,000 rpm for 10 min, and the supernatants were used for preparation of culture media. Data represent means ± SD of 3 or 13 independent experiments. *p < 0.05; **p < 0.01.

Regarding the NaHCO_3_ concentration, the effect of NaHCO_3_ was significant at 1 mg/ml and subsequently increased in a dose-dependent manner up to 5 mg/ml. Thereafter, the effect of NaHCO_3_ gradually decreased in the range of 7–10 mg/ml (Fig. 2A). For Na_2_CO_3_, a significant effect was observed at 0.5 mg/ml and the maximum effect (~200% of control) was observed at 2 mg/ml (Fig. 2B). The latter concentration is close to the serum bicarbonate concentrations (2.0–2.4 mg/ml) that improved serum glucose and insulin levels in diabetic chronic kidney disease patients with acidosis after oral administration of bicarbonate *in vivo* [30]. The effect of Na_2_CO_3_ decreased rapidly in the range of 4–5 mg/ml, and completely inhibited glucose consumption by cultured cells at >7 mg/ml. Both carbonates accelerated glucose consumption in TE-13 cells derived from human esophageal cancers [32] (data not shown).

**Figure 2.**
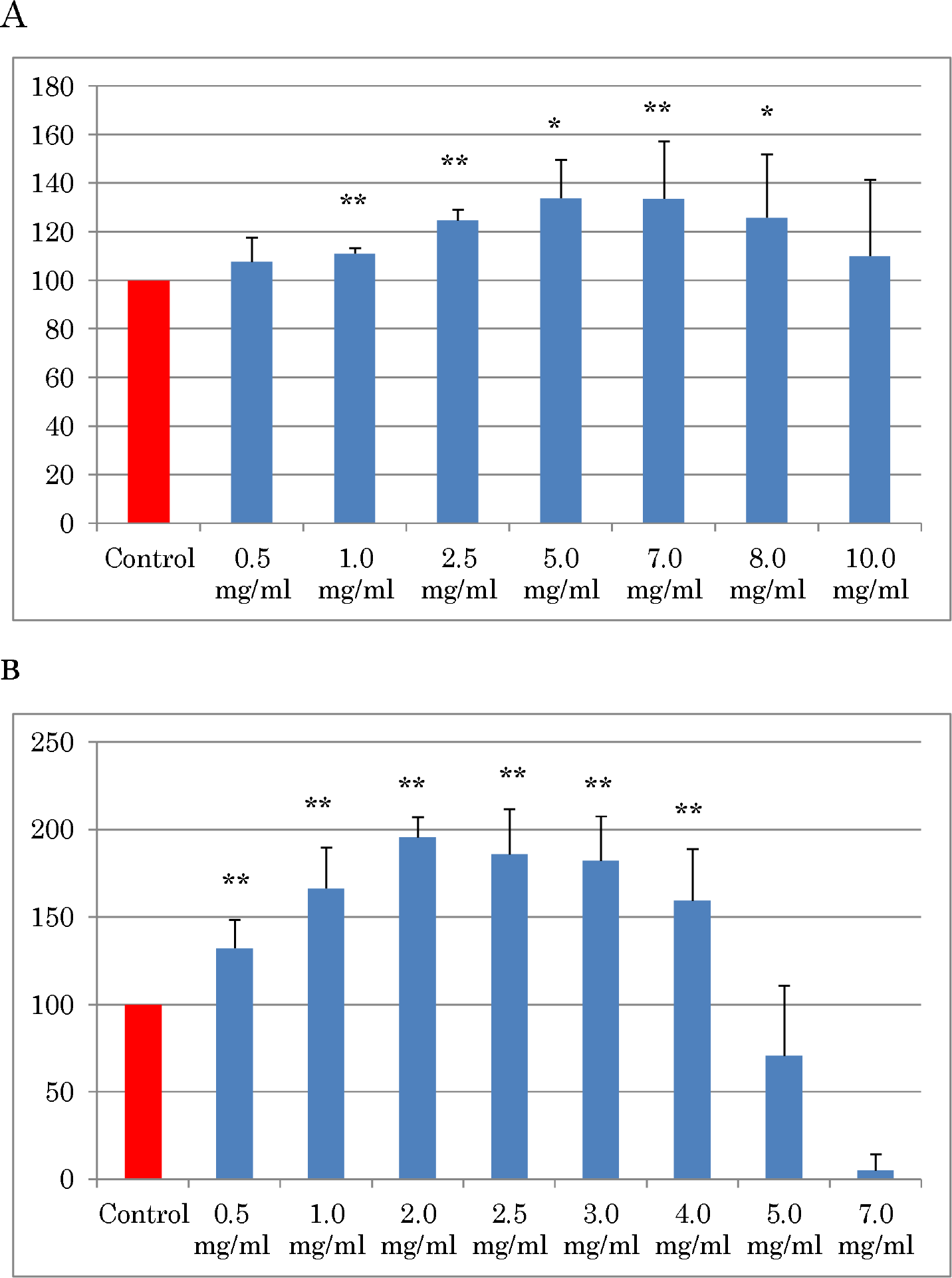
Effect of NaHCO_3_ (A) or Na_2_CO_3_ (B) on glucose consumption by Py-3Y1-S2 cells. Data represent means ± SD of 3–7 independent experiments. *p < 0.05; **p < 0.01.

### Cellular proteins

NaHCO_3_ and Na_2_CO_3_ solutions are alkaline, and their pH values in culture media were 8.0–9.5 in the presence of 5% CO_2_ in the cell incubator. Of course, culture medium initially contains NaHCO_3_ to maintain the pH at around 7.5. The cellular protein contents of Py-3Y1-S2 cells were independent of NaHCO_3_ in the range of 0.5–8 mg/ml, while NaHCO_3_ at 10 mg/ml reduced the cellular protein to 70% of the control (Fig. 3). The decrease in cellular protein content was linked with the disappearance of glucose consumption acceleration at the high concentration of NaHCO_3_ (Fig. 2A). Similarly, the Py-3Y1-S2 cellular protein content was markedly reduced by 5 mg/ml Na_2_CO_3_ to around 30% of the control (Fig. 2B). The cellular protein content was reduced to <10% in the presence of >7 mg/ml Na_2_CO_3_. This cytotoxic effect of Na_2_CO_3_ was in accordance with the decrease in glucose consumption (Fig. 3), and was caused by the high pH of 9.5 in the absence of CO_2_. The culture medium containing 5 mg/ml Na_2_CO_3_ had a pH of ~8.5 after incubation in the presence of 5% CO_2_. Therefore, conditions involving high pH above 9 led to cell death.

**Figure 3.**
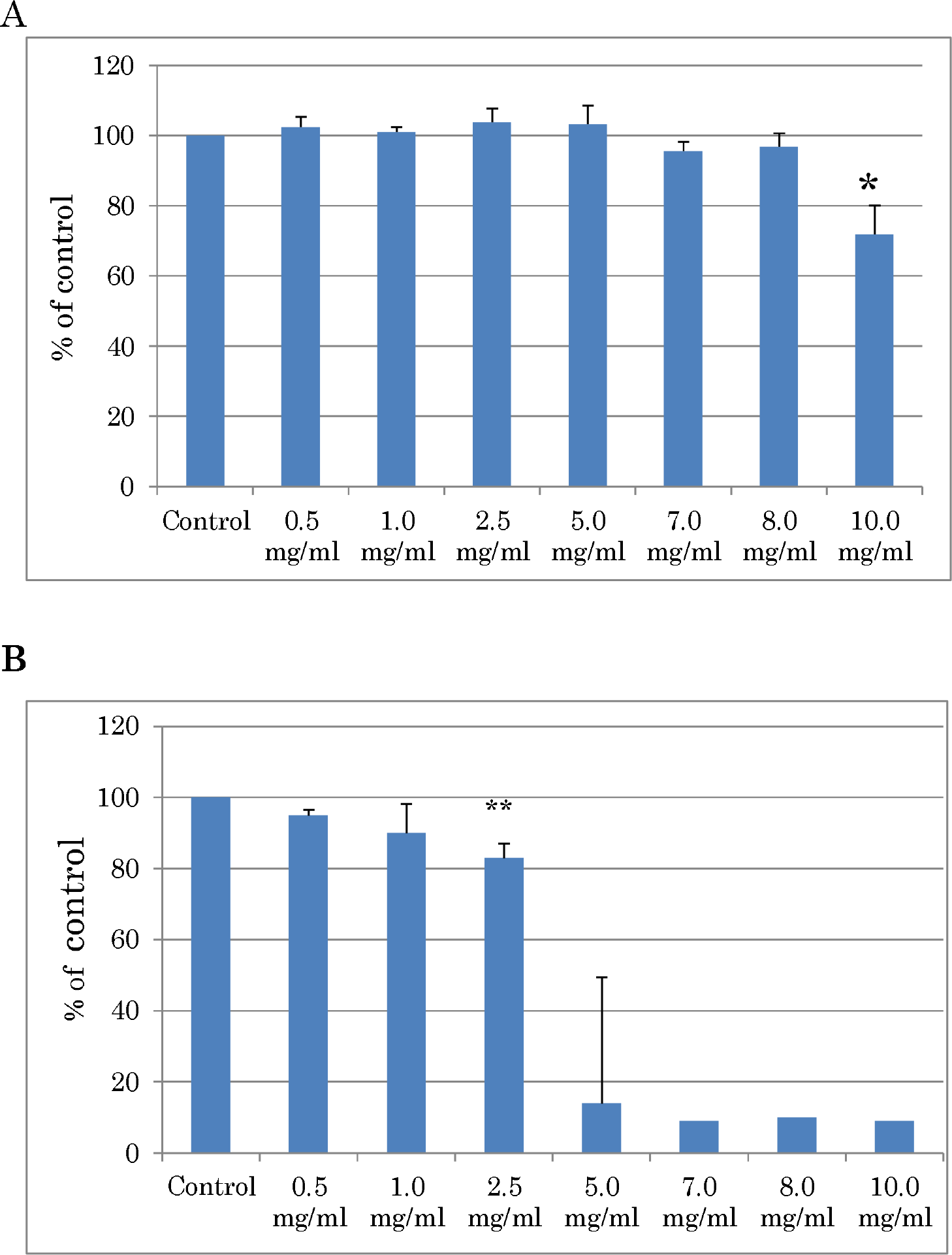
Effect of NaHCO_3_ (A) or Na_2_CO_3_ (B) on cellular protein. Data represent means ± SD of 5 independent experiments. *p < 0.05; **p < 0.01.

### Additive effects of Con A or vanadium

The lectin Con A was reported to exert insulin-like effects during *in vitro* experiments [33]. We showed that Con A shared insulin receptors with insulin by *in vitro* experiments involving binding assays of labeled Con A and insulin [34,35]. When cultured Py-3Y1-S2 cells were treated with Con A at 0.025 mg/ml, glucose consumption increased in a dose-dependent manner (Fig. 4). Con A at 0.1 mg/ml further increased glucose consumption, and the effect almost reached a plateau. Thus, the effects of Con A at 0.1 mg/ml were examined at different NaHCO_3_ concentrations. NaHCO_3_ at 0.5 mg/ml and 1.0 mg/ml further increased glucose consumption with Con A at 0.1 mg/ml, while the effect of NaHCO_3_ at 5.0 mg/ml was almost null in the presence of 0.1 mg/ml Con A (Fig. 5A). Similarly, Na_2_CO_3_ at 0.5 mg/ml further increased glucose consumption with Con A at 0.1 mg/ml (Fig. 5B). These results indicate that combinations of NaHCO_3_ or Na_2_CO_3_ with Con A show sufficient effects even low concentrations compared with their use alone.

**Figure 4.**
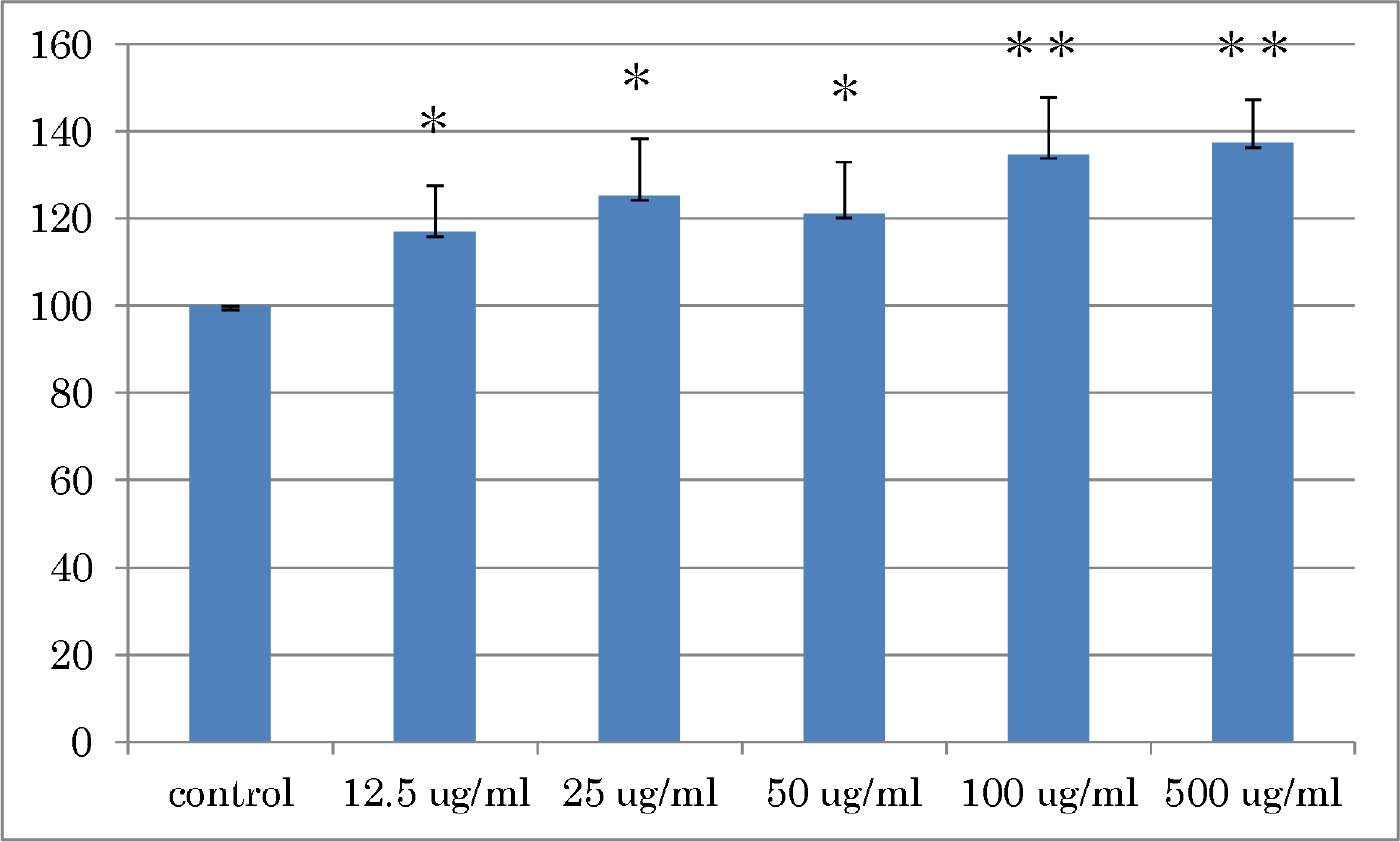
Effect of Con A on glucose consumption by Py-3Y1-S2 cells. Data represent means ± SD of 5 independent experiments. *p < 0.05; **p < 0.01.

**Figure 5.**
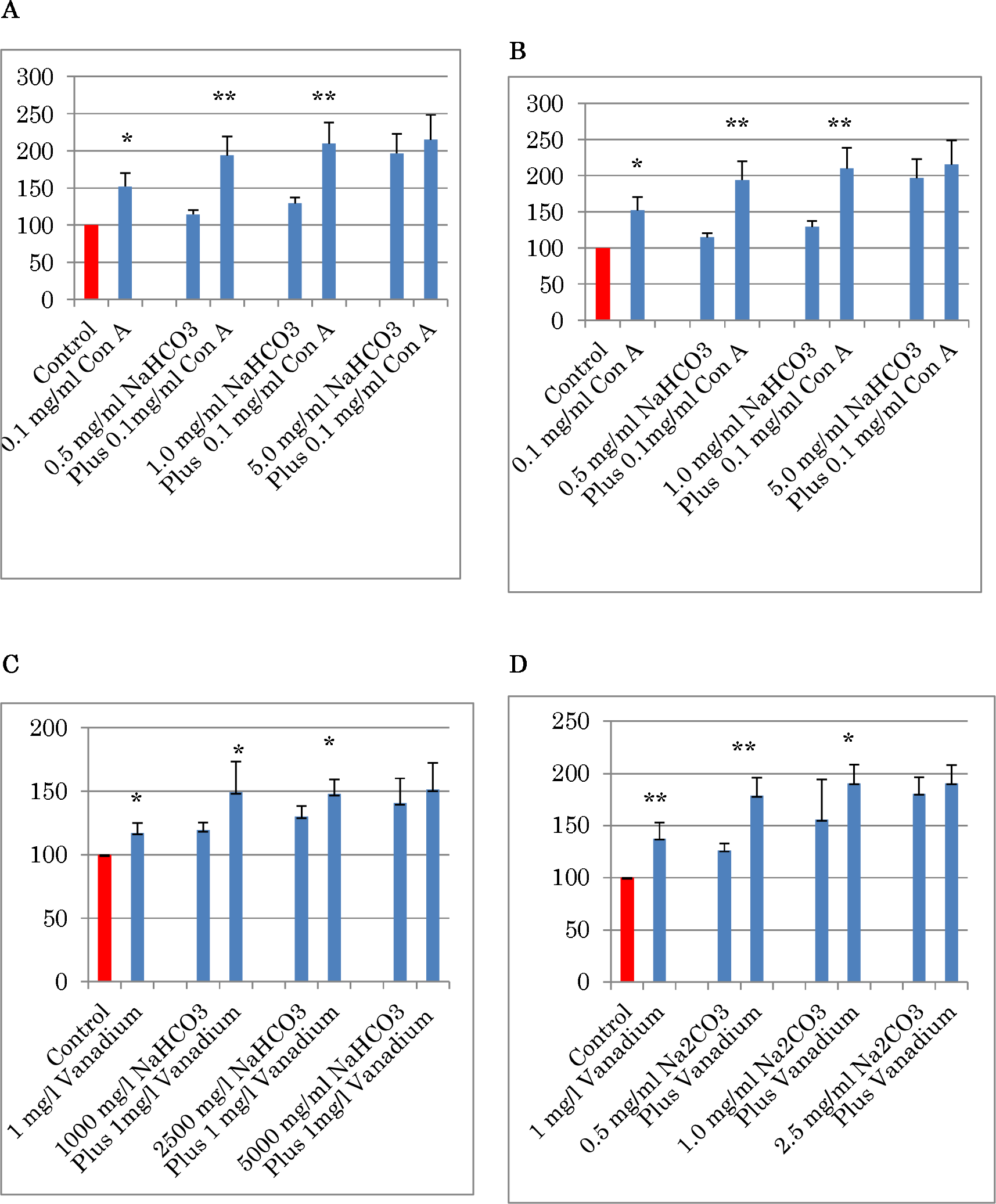
Effect of vanadium or Con A on glucose consumption by Py-3Y1-S2 cells in the presence of NaHCO_3_ (A and B) or Na_2_CO_3_ (C and D). Data represent means ± SD of 5 independent experiments. *p < 0.05; **p < 0.01.

In a previous study [24], vanadium accelerated glucose consumption in a dose-dependent manner in the range of 25–10,000 μg/L, while an inhibitory effect was observed at >1.0 mg/L in Py-3Y1-S2 cells. When cells were cultured with 1.0 mg/L vanadium plus 1.0, 2.5, or 5.0 mg/ml NaHCO_3_, glucose consumption was significantly accelerated in the presence of 1.0 and 2.5 mg/ml NaHCO_3_ (Fig. 5C). However, no additive effect of vanadium was observed at 5.0 mg/ml NaHCO_3_. Similarly, lower concentrations of Na_2_CO_3_, 0.5 mg/ml and 1.0 mg/ml, exhibited a significant additive effect with vanadium, while 5.0 mg/ml Na_2_CO_3_ had no additive effect with vanadium (Fig. 5D).

### Effects of diabetic reagents

STZ, alloxan, and nicotinamide are known chemical compounds that can induce diabetes in animals [36,37]. In animals, alloxan destroys ß-cells in the pancreas at high concentrations, and induces type 2 diabetes at low concentrations. When Py-3Y1-S2 cells were cultured in the presence of the above drugs *in vitro*, the glucose consumption in the culture medium was significantly reduced (Fig. 6A). In the present study, the drugs did not cause significant reductions in cellular protein amounts (data not shown). In contrast, addition of NaHCO_3_ or Na_2_CO_3_ to the culture medium abolished the effects of STZ, alloxan, and nicotinamide in Py-3Y1-S2 cells (Fig. 6B). These findings indicate that the reduction in glucose consumption by Py-3Y1-S2 cells was not caused by cell death.

**Figure 6.**
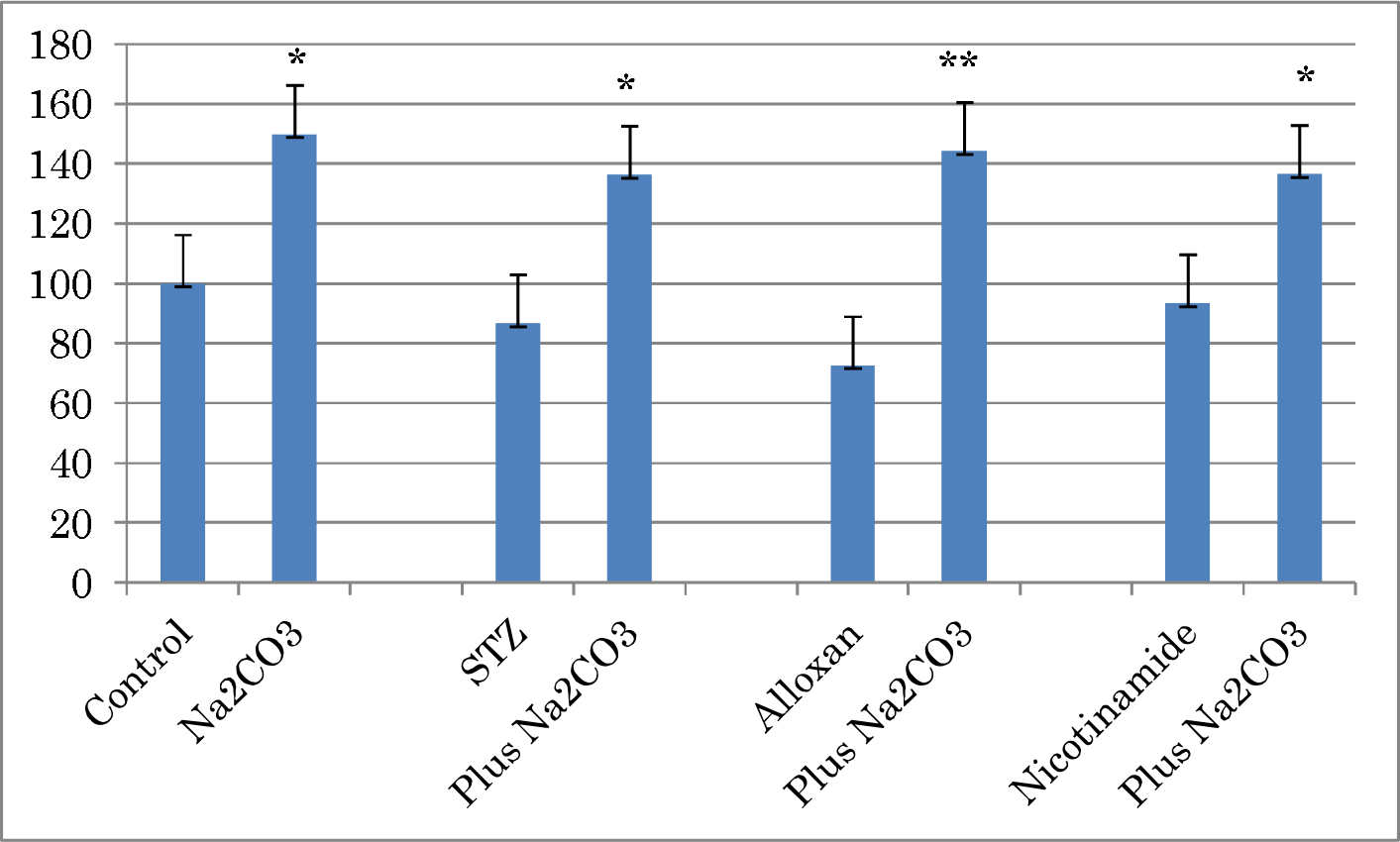
Effect of NaHCO_3_ (A) or Na_2_CO_3_ (B) on glucose consumption by Py-3Y1-S2 cells in the presence of STZ, alloxan, or nicotinamide. The concentration of each reagent was 1 mg/ml. Data represent means ± SD of 3 independent experiments. *p < 0.05; **p < 0.01.

Glucose is metabolized to pyruvic acid during the first stage in the absence of oxygen, and then alcohols or organic acids are formed via the TCA cycle based on various enzyme reactions in cells. During the second stage in the presence of oxygen, pyruvic acid is finally converted to CO_2_ and H_2_O. Thus, NaHCO_3_ or Na_2_CO_3_ formed from CO_2_ could be involved in glucose regulation like insulin. These regulatory mechanisms by carbonates resemble the autocrine or paracrine mechanisms of cytokines. When PY-3Y1-S2 cells were cultured in the presence of NaHCO_3_ or Na_2_CO_3_, the concentration of lactate in the culture medium was significantly increased (Fig. 7). Meanwhile, STZ, alloxan, and nicotinamide significantly reduced lactate production, and further addition of NaHCO_3_ or Na_2_CO_3_ abolished the effects of these drugs.

**Figure 7.**
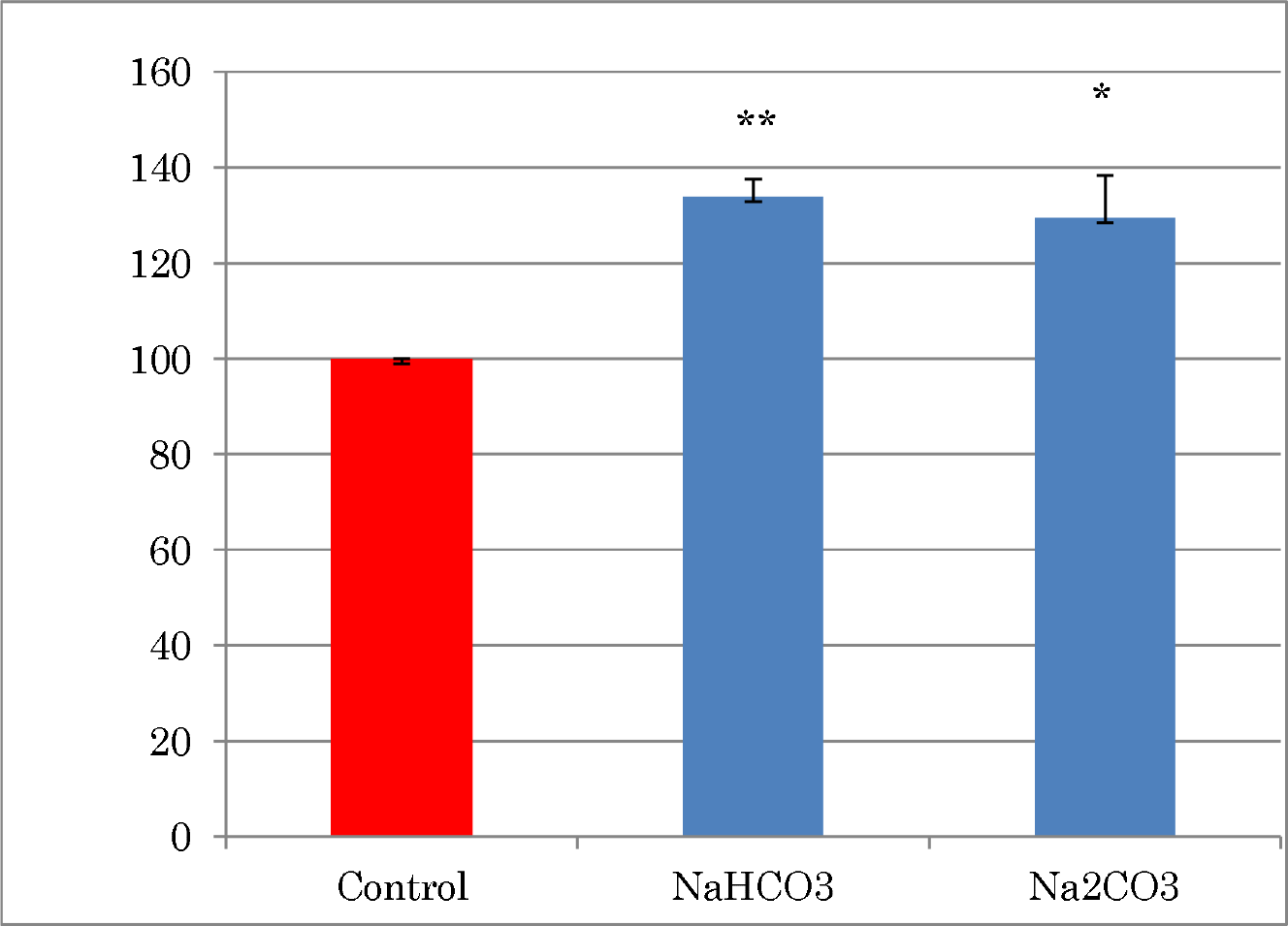
Effect of NaHCO_3_ or Na_2_CO_3_ on lactate production by Py-3Y1-S2 cells. Data represent means ± SD of 6–8 independent experiments. *p < 0.05; **; p < 0.01.

The basic structure of both NaHCO_3_ and Na_2_CO_3_, which accelerate glucose consumption, is a carbonyl group, =C=O, while the structures of STZ, alloxan, and nicotinamide, which induce type 1 diabetes and reduce glucose consumption in cultured cells, also contain a carbonyl group. The carbonyl group of diabetes-inducing reagents like STZ, alloxan, and nicotinamide is a part of an amide structure, -CONHR. Ceramides consisting of an amino group-bound carbonyl group were shown to induce insulin resistance [38]. However, amino acids containing a carboxyl group did not affect glucose uptake in the present study, although their structures contained a carbonyl group that was not bound to an amino group. These findings indicate that glucose consumption by NaHCO_3_ or Na_2_CO_3_ is controlled by side groups that bind to the carbonyl group, such that the –O^-^ accelerates glucose consumption and the –N^-^ reduces glucose consumption.

NaHCO_3_ is used as baking soda in daily life and as a medicine for alkaloids. However, because NaHCO_3_ and Na_2_CO_3_ are alkaline chemical compounds, their concentrations for usage should be carefully controlled. The present results may contribute to the development of alternative medicines as well as new medicines for DM and dementia.

### Lactate production

When Py-3Y1-S2 cells were cultured in DMEM, lactate was secreted into the culture medium (Fig. 7). Further addition of NaHCO_3_ or Na_2_CO_3_ to the culture medium accelerated lactate secretion by the cells. These findings indicate that carbonates directly regulate glucose metabolism in Py-3Y1-S2 cells.

## CONCLUSIONS

Two carbonates, NaHCO_3_ and Na_2_CO_3_, directly accelerated glucose consumption in established Py-3Y1-S2 rat fibroblast cells. Combinations of the carbonates with vanadium or Con A further increased glucose consumption, while combinations of the carbonates with STZ, alloxan, or nicotinamide abolished the effects of the drugs in Py-3Y1-S2 cells. Both carbonates increased lactate production by Py-3Y1-S2 cells. Taken together, the present study indicates that NaHCO_3_ and Na_2_CO_3_ directly regulate glucose metabolism in Py-3Y1-S2 cells.

## Acknowledgments

The author thanks Mr. Fumio Takei of Fuji Yogan Institute for his kind donation of Mt. Fuji basal powder. The author also thanks Alison Sherwin, PhD, from Edanz Group (www.edanzediting.com/ac) for editing a draft of this manuscript.

